# Genetic variation modulates susceptibility to aberrant DNA hypomethylation and imprint deregulation in naïve pluripotent stem cells

**DOI:** 10.1101/2024.06.26.600805

**Authors:** C Parikh, RA Glenn, Y Shi, K Chatterjee, EE Swanzey, S Singer, SC Do, Y Zhan, Y Furuta, M Tahiliani, E Apostolou, A Polyzos, R Koche, JG Mezey, T Vierbuchen, M Stadtfeld

## Abstract

Naïve pluripotent stem cells (nPSC) frequently undergo pathological and not readily reversible loss of DNA methylation marks at imprinted gene loci. This abnormality poses a hurdle for using pluripotent cell lines in biomedical applications and underscores the need to identify the causes of imprint instability in these cells. We show that nPSCs from inbred mouse strains exhibit pronounced strain-specific susceptibility to locus-specific deregulation of imprinting marks during reprogramming to pluripotency and upon culture with MAP kinase inhibitors, a common approach to maintain naïve pluripotency. Analysis of genetically highly diverse nPSCs from the Diversity Outbred (DO) stock confirms that genetic variation is a major determinant of epigenome stability in pluripotent cells. We leverage the variable DNA hypomethylation in DO lines to identify several trans-acting quantitative trait loci (QTLs) that determine epigenome stability at either specific target loci or genome-wide. Candidate factors encoded by two multi-target QTLs on chromosomes 4 and 17 suggest specific transcriptional regulators that contribute to DNA methylation maintenance in nPSCs. We propose that genetic variants represent candidate biomarkers to identify pluripotent cell lines with desirable properties and might serve as entry points for the targeted engineering of nPSCs with stable epigenomes.

**Highlights:**

- Naïve pluripotent stem cells from distinct inbred mouse strains exhibit variable DNA methylation levels at imprinted gene loci.
- The vulnerability of pluripotent stem cells to loss of genomic imprinting caused by MAP kinase inhibition strongly differs between inbred mouse strains.
- Genetically diverse pluripotent stem cell lines from Diversity Outbred mouse stock allow the identification of quantitative trait loci controlling DNA methylation stability.
- Genetic variants may serve as biomarkers to identify naïve pluripotent stem cell lines that are epigenetically stable in specific culture conditions.

## Introduction

Naïve pluripotent stem cells (nPSCs) resembling the pre-implantation embryo are capable of extensive *ex vivo* self-renewal, amenable to genome engineering, and can differentiate into all somatic cell types. This combination of features makes nPSCs, in principle, tailor-made for regenerative medicine applications. However, several molecular abnormalities manifesting in cultured pluripotent cells complicate exploiting this potential (Andrews et al., 2022). For example, the pervasive epigenetic instability of nPSCs upon *ex vivo* culture – which manifests as aberrant changes in DNA methylation and other chromatin marks that compromise physiological transcriptional programs and functional properties (Mani and Mainigi, 2018; Meissner et al., 2008; Rebuzzini et al., 2016) – represents a significant roadblock for many biomedical applications of PSCs.

Aberrant DNA methylation changes in cultured nPSCs include hypermethylation and hypomethylation events (Habibi et al., 2013; Lee et al., 2018) and affect gene loci encoding essential developmental regulators. Solidifying naïve pluripotency by chemical inhibition of mitogen-activated protein kinase (MAPK) signaling, an approach often applied for both mouse (Ficz et al., 2013; Leitch et al., 2013) and human cells (Bayerl et al., 2021; Pastor et al., 2016), results in widespread loss of DNA methylation in both species. Epigenetic changes are particularly problematic at imprinted genes since the loss of the parent-of-origin asymmetry in DNA methylation at these loci cannot be readily restored (Bayerl et al., 2021; Pastor et al., 2016) and is associated with specific defects in embryonic development (Ferguson-Smith and Bourc’his, 2018) that complicate disease modeling with affected cells

DNA methylation abnormalities in naïve nPSCs are associated with a high degree of line-to-line variability, even among cell lines established under identical conditions (Bock et al., 2011; Humpherys et al., 2001; Johannesson et al., 2014; Lin and Xiao, 2017). This has given rise to the notion that randomly occurring pathological epigenome changes in nPSCs are an unavoidable side-effect of the extraordinary developmental flexibility of these cells. Several recent studies leveraging distinct inbred mouse strains with fully sequenced genomes (Lilue et al., 2018) and high-resolution panels of single nucleotide polymorphisms (SNPs)(Morgan et al., 2015) have shown that core PSC properties such as self-renewal capacity (Skelly et al., 2020) and *in vitro* differentiation bias (Byers et al., 2022; Ortmann et al., 2020) are modulated by genetic variation. In addition, pathological DNA hypermethylation at the *Dlk1-Dio3* locus in nPSCs is controlled by a trans-acting quantitative trait locus (QTL) that distinguishes the commonly used B6J and 129 strains (Swanzey et al., 2020). Whether this observation extends to other imprinted gene loci and, most importantly, to the biomedically highly relevant susceptibility of nPSCs for DNA hypomethylation remains unanswered.

Here, we use nPSCs from a combination of distinct inbred strains and a genetically diverse outbred stock (Diversity Outbred; DO mice) to systematically characterize the impact of genetic variation on loss of DNA methylation at ICRs. Our data indicate that susceptibility to DNA hypomethylation in nPSCs is determined by identifiable genetic variants, reveal candidate regulators of DNA methylation levels via QTL mapping, and suggest rational approaches to stabilize the epigenome of naïve pluripotent stem cells.

## Results

### Strain-specific introduction of imprint abnormalities in iPSCs established from inbred mice

To begin investigating the degree to which genetic background influences the stability of imprinting marks during nPSC derivation and maintenance, we used OKSM reprogramming (Sommer et al., 2009) to establish induced pluripotent stem cells (iPSCs) from mouse embryonic fibroblast (MEFs) derived from seven distinct inbred mouse strains (129S1/SVlmJ; 129, C57BL6/J; B6J, C57BL6/NJ; B6N, CBA/J; CBA, DBA/2J; DBA, C3H/HeJ; C3H, and A/J; AJ) (**Fig. S1A**). We cultured cells undergoing reprogramming in media containing ascorbic acid (AA) and modulators of WNT and TGFβ signaling (**Fig. 1A**), which dramatically facilitates iPSC formation (Vidal et al., 2014). Since AA has been shown to stimulate TET enzymes (Blaschke et al., 2013; Monfort and Wutz, 2013), we reasoned that transient exposure to this compound might also reveal strain-specific susceptibilities to DNA demethylation. In contrast, subsequent culture in standard serum-containing media (**Fig. 1A**) would reveal susceptibilities to DNA hypermethylation, as we have previously shown for *Dlk1-Dio3* (Swanzey et al., 2020). For these experiments, we focused exclusively on male cells to avoid confounding the effects of genetic variants with the well-documented DNA hypomethylation propensity observed in female nPSCs due to the presence of two active X chromosomes (Zvetkova et al., 2005). We obtained stable iPSC colonies independent of transgenic OKSM expression from each of the seven inbred strains, including the CBA and DBA strains that are not permissive for ESC derivation in standard conditions (Czechanski et al., 2014), albeit at variable efficiencies (**Fig.S1B**). Expanded polyclonal iPSC lines (n=2 biological replicates from each inbred strain; **Table S1**) exhibited the expected pluripotent cell morphology (**Fig. 1B**) and expression of the pluripotency-associated markers SSEA1 and EpCAM (**Fig. S1C**) (Polo et al., 2012).

**Figure 1.**
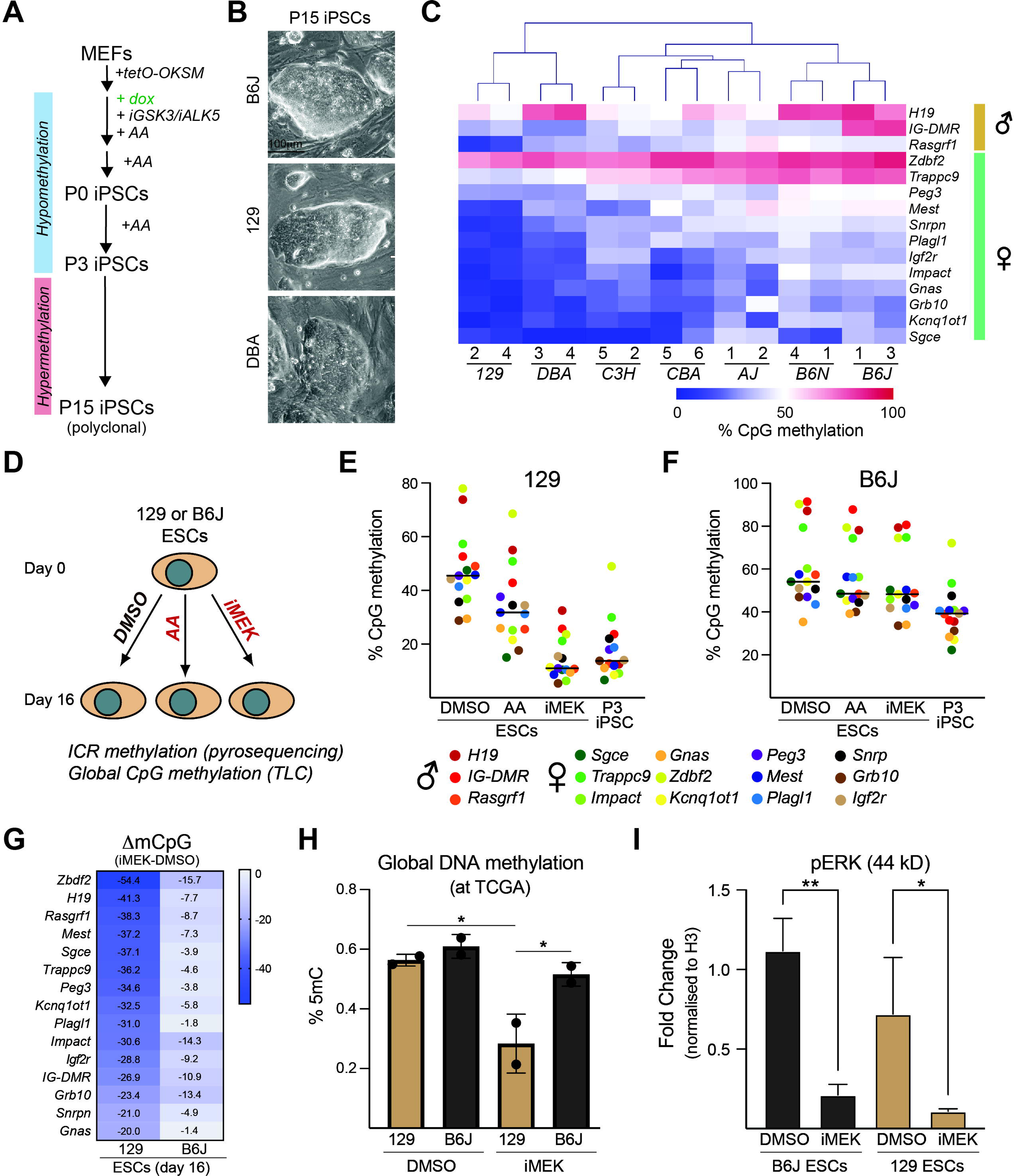
Genetic variation modulates DNA hypomethylation at ICRs in naive pluripotent stem cells from distinct inbred mouse strains. A. Experimental strategy to derive polyclonal iPSCs from MEFs derived from distinct inbred strains. B. Representative colony morphology of iPSCs from each inbred strain at passage 15 (p15). C. Unsupervised clustering of CpG methylation levels at selected ICRs in polyclonal iPSCs (n=2 biological replicates/strain). D. Strategy to test the impact of AA or iMEK on imprint methylation in 129 and B6J mESCs. TLC = thin layer chromatography. E. DNA methylation levels at ICRs in indicated cell types derived from the 129 strain F. Same as (E) but for the B6J strain. G. Average changes in DNA methylation levels (iMEK minus DMSO) at germline ICRs in 129 and B6J ESCs. H. Quantification of global DNA methylation levels at TCGA sequences by thin layer chromatography in indicated mESCs after 16 days of culture in either the presence of DMSO or of iMEK. * = p<0.05 with one-way ANOVA. N = 2 biological repeats. I. Quantification of phosphorylated-ERK protein levels in 129 and B6J mESCs after culture in iMEK or DMSO. Protein levels are normalized to histone H3 expression. * = p<0.05. and ** = p<0.01 with one-way ANOVA. N = 3 biological replicates.

To assess the status of imprinted gene regulation, we subjected genomic DNA (gDNA) isolated from passage 15 (P15) iPSCs from each inbred strain for DNA methylation analysis by targeted bisulfite sequencing. We analyzed established control regions of the three paternally imprinted loci (*Dlk1-Dio3*, *H19/Igf2*, *Rasgrf1*) and twelve maternally imprinted loci (*Gnas*, *Grb10*, *Igf2r*, *Impact*, *Inpp5f*, *Kcnq1ot1*, *Mest*, *Nap1l5*, *Nnat*, *Peg3*, *Plagl1*, *Sgce*, *Snrpn*, *Trappc9*, *Zdbf2* and *Zrsr1*). Unsupervised clustering of these DNA methylation data revealed considerable locus-to-locus and strain-to-strain variability in DNA methylation levels, with biological replicates from each genetic background clustering together (**Fig. 1C**). This suggests a significant contribution of genetic background to imprinted gene stability. For further analyses, we defined hypermethylation as >70% CpG methylation and hypomethylation as <20% CpG methylation at a given locus based on average values from biological replicates. With these criteria, hypermethylation of *Dlk1-Dio3 (*controlled by the intergenic differentially methylated region, short *IG-DMR)* was only observed in B6J-iPSCs but not in iPSCs from the other inbred strains, including the closely related B6N strain (**Fig. S1A**). These data suggest that the genetic variant(s) predisposing nPSCs to pathological DNA hypermethylation (Swanzey et al., 2020) are uniquely present or active in B6J mice. Additional evidence that genetic background controls locus-specific DNA hypermethylation comes from analysis of *H19* (affected in DBA-, B6N- and B6J-iPSCs) and *Trappc9* (affected in C3H-, CBA-, AJ-, B6N- and B6J-iPSCs) (**Fig. 1C**). In contrast, *Zdbf2* exhibited hypermethylation in iPSCs from all inbred strains, suggesting that this locus might be particularly susceptible to acquire hypermethylation in iPSCs (**Fig. 1C**). Focusing on DNA hypomethylation, we observed multi-locus loss of imprint methylation in iPSCs from 129 (10 loci), DBA (7 loci), CBA (4 loci) and C3H (2 loci) (**Fig. 1C**), indicating either the presence of multiple variants that affect individual genes or the existence of variants that affect DNA methylation at multiple loci.

To further investigate strain-specific susceptibilities to DNA methylation change and to determine when, during the iPSC derivation process, they might manifest, we focused on 129 and B6J, the two backgrounds exhibiting the most divergent ICR DNA methylation profiles in P15 iPSCs (**Fig. 1C**). While MEFs from 129 and B6J mice both showed indistinguishable, physiological DNA methylation levels at all ICRs (**Fig. S1D**), P3 129 iPSCs (methylation analysis concomitant with AA withdrawal; see **Fig. 1A**) already exhibited pronounced DNA hypomethylation likely introduced during the reprogramming process. In contrast, DNA hypermethylation was not evident at ICRs in B6J iPSCs at this earlier stage of derivation (**Fig. S1D**), suggesting DNA hypermethylation is introduced during prolonged culture in the absence of AA. Together, these observations reveal strain-specific differences in the stability of DNA methylation levels at imprinted gene control regions in iPSCs derived under identical conditions specifically selected to reveal both susceptibilities to DNA hypo- and hypermethylation.

### Susceptibility to DNA hypomethylation upon MAP kinase inhibition is governed by genetic background

To determine whether nPSCs derived using other standard approaches also exhibit strain-specific differences to loss-of-imprinting by DNA hypomethylation as observed in iPSCs, we exposed ESCs established from 129 and B6J blastocysts to either AA or the MAPK inhibitor PD0325901 (“iMEK”) for 16 days (**Fig. 1D**), resembling the time required to reprogram MEFs into early-stage iPSCs. 129 ESCs cultured in the presence of AA showed evidence for DNA hypomethylation, albeit significantly less severe than P3 iPSCs (**Fig. 1E** and **Fig.S1E**), suggesting that the reprogramming process can exacerbate the loss of methylation at ICRs. In contrast, 129 ESCs exposed to iMEK showed dramatic loss of ICR methylation at all loci studied, similar to what was observed in P3 iPSCs (**Fig. 1E**) and in agreement with the reported role of MEK inhibition in DNA demethylation (Choi et al., 2017; Yagi et al., 2017). Compared to 129 ESCs, B6J ESCs cultured with either AA or iMEK exhibited a modest reduction in ICR methylation, which remained in the physiological range (**Fig. 1F** and **Fig.S1E**). This documents a surprising degree of resistance of B6J ESCs to pathological DNA hypomethylation that extends to all ICRs analyzed (**Fig. 1G**). A somewhat less pronounced resistance to loss of ICR methylation was observed in B6J:129 F1 hybrid ESCs (**Fig. S1F**).

We conducted thin-layer chromatography experiments to determine whether loss of ICR methylation in 129 mESCs reflects genome-wide changes in total DNA methylation levels. This approach revealed significantly reduced global levels of DNA methylation at CpG residues in 129 mESCs but not in B6J exposed to iMEK. In contrast, mESCs from both strains that were cultured in standard conditions showed similar DNA methylation levels (**Fig. 1H**). These observations suggest that the effects of genetic background on susceptibility to DNA hypomethylation downstream of MAPK inhibition are not restricted to ICRs, which is consistent with prior observations made in mouse and human nPSCs (Choi et al., 2017; Pastor et al., 2016). Of note, the reduction in ERK phosphorylation (**Fig. 1I**) and the levels of DNMT3A (**Fig. S1G**) were similar in 129 and B6J ESCs exposed to iMEK. These results demonstrate that the observed differences in DNA methylation stability between mESCs from these two strains are not due to the different effectiveness of the inhibitor. They also suggest that the responsible variants influence the recruitment or activity of DNA methyltransferases rather than directly altering the expression levels of these enzymes. Together, our findings demonstrate that susceptibility to pathological loss of DNA methylation in commonly used nPSC culture conditions is strongly modulated by genetic variation.

### Variable DNA hypomethylation in naïve PSCs derived from a genetically diverse outbred mouse stock

Our results so far document that genetic variation between nPSCs from different inbred mouse strains results in markedly different susceptibility to hypomethylation of ICRs driven by MAPK inhibition. We sought to determine whether a similar effect is evident in genetically diverse nPSCs that more closely represent the human population and would potentially allow mapping underlying variants. To this end, we used a pooled *in vitro* fertilization (IVF) approach to establish a panel of ESC lines from Diversity Outbred (DO) mice (Glenn et al., 2024) (**Fig. 2A**), a heterogenous outbred stock derived from eight founder strains (A/J, C57Bl/6J [B6J], 129S1/SvImJ [129], NOD/ShiLtJ [NOD], NZO/HlLtJ [NZO], CAST/EiJ [CAST], PWK/PhJ [PWK] and WSB/EiJ [WSB]) that harbors more than 40 million SNPs and structural variants and allows high-resolution genetic mapping (Gatti et al., 2014; Melia and Waxman, 2020; Smallwood et al., 2014; Swanzey et al., 2021). For our experiments, we collected genomic DNA from a panel of 85 male DO mESC lines that exhibited undifferentiated morphology and were cultured in the presence of iMEK for 2-4 passages to promote hypomethylation (**Fig. 2A**). Kinship analysis after SNP genotyping confirmed a low degree of genetic relatedness among these lines (**Fig.S2A**).

**Figure 2.**
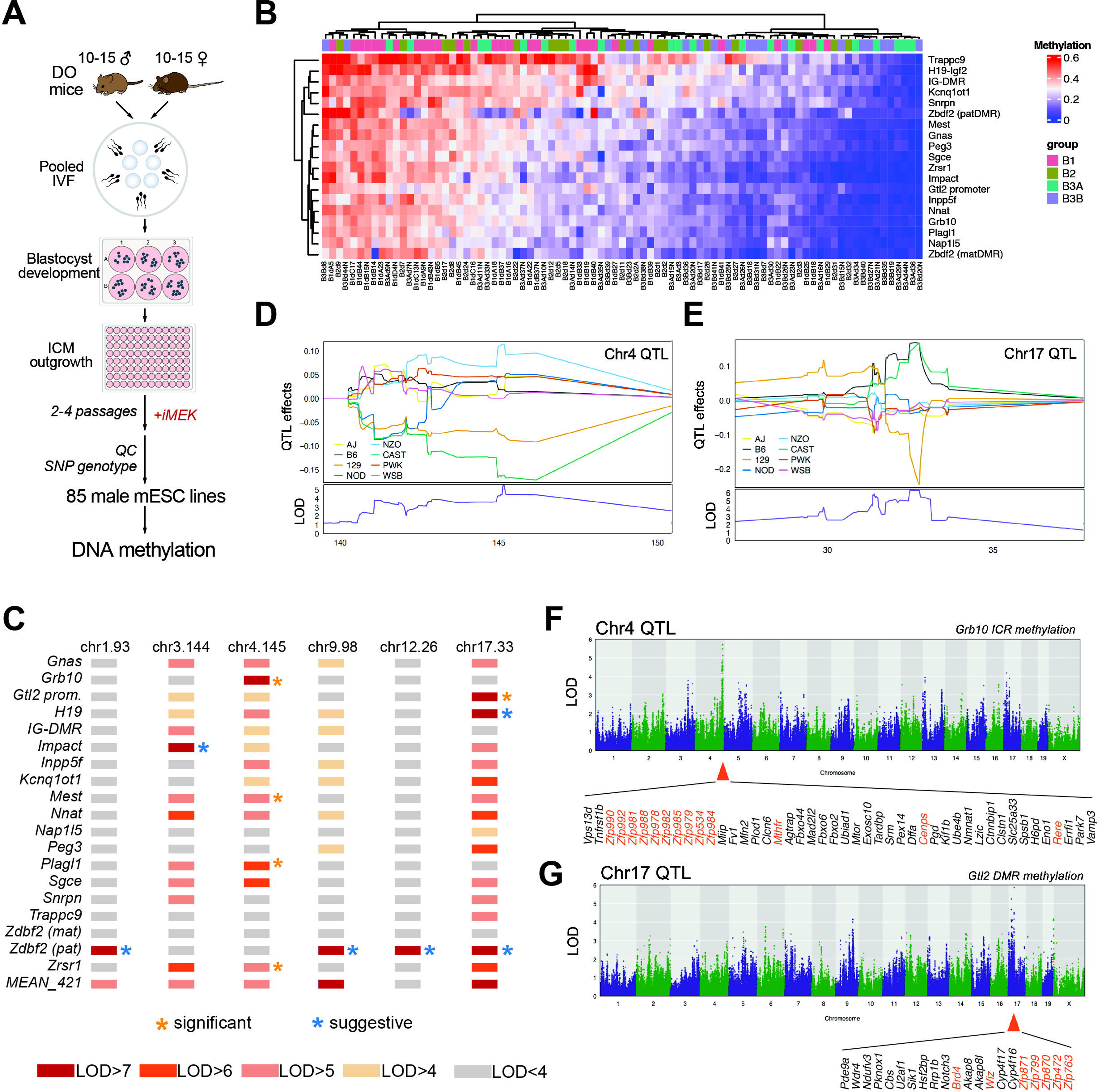
Identification of QTLs modulating pathological DNA hypomethylation at ICRs in naïve PSCs. A. Overview of experimental strategy. mESCs from DO mice were generated via a pooled in vitro fertilization approach. DO mESC lines (n = 85 male lines) were banked at early passage (p3) and then thawed and cultured for an additional two passages in serum/LIF + 2i to promote hypomethylation of ICRs. B. ICR DNA methylation levels across all of the DO mESC lines. C. Genome-wide significant (FWER<0.1) and suggestive (FWER<0.2) QTLs controlling ICR methylation across the mouse genome. D. Allelic effect plot of the Chr4 QTL. E. Allelic effect plot of the Chr17 QTL. F. Genome-wide plot of LOD scores for methylation at the *Grb10* ICR with the Chr4 QTL indicated. Shown are encoded genes with tpm>10 in either B6J or 129 mESCs, with known transcriptional or chromatin regulators highlighted in orange. G. Genome-wide plot of LOD scores for methylation at the *Gtl2* promoter with the Chr17 QTL indicated. Show are encoded genes with tpm>10 in either B6J or 129 mESCs, , with known known transcriptional or chromatin regulators highlighted in orange.

To sensitively measure DNA methylation levels at ICRs and at additional selected loci across the genome (**Fig.S2B**)(**Table S2**) in DO ESC lines, we used Enzymatic Methyl (EM)-seq and a custom targeted capture panel. Unbiased clustering of DNA methylation levels at all regions revealed markedly different DNA methylation levels across the panel of DO nPSC lines (**Fig.S2C**). ICRs had the highest average levels of DNA methylation among the different categories of cis-regulatory elements included within the targeted capture panel (**Fig.S2D**). Importantly, differences in CpG methylation levels were not correlated with differences in sequencing coverage (**Fig.S2E**), confirming the reliability of our approach. These data suggest that ICRs in nPSCs might have increased protection from demethylation compared to other gene loci, potentially reflecting their increased resistance to genome-wide epigenetic reprogramming in the pre-implantation embryo (Monk, 2015). Nevertheless, DNA methylation levels at ICRs were also highly variable between DO cell lines, ranging from physiological levels (∼50%) of DNA methylation at all ICRs in some cell lines to essentially complete loss of methylation at all ICRs in other lines (**Fig.2B**). These data suggest that genetic variation present within the DO stock can influence DNA hypomethylation phenotypes in naïve PSCs cultured with iMEK.

To evaluate whether different ICRs respond similarly to MAPK inhibition, we performed principal component analysis (PCA) on ICR DNA methylation data across the panel of DO nPSC lines. This analysis indicated that most ICRs respond in a strongly correlated manner (**Fig.S2F**), except for 1) regions controlling imprinting at the two major paternally imprinted gene clusters (IG-DMR and *H19*-*Igf2*), 2) a somatic DMR at the maternally imprinted *Zdbf2* locus that acquires DNA methylation on the paternal allele late during embryogenesis (Duffie et al., 2014) and C) *Trappc9*, a locus encoding transcripts known to exhibit variable patterns of parent-of-origin specific imprinting during brain development (Claxton et al., 2022). Consistent with our results with iPSCs from distinct inbred strains, these data provide extensive additional evidence that the majority of ICRs undergo pathological loss of DNA methylation in response to iMEK treatment and that genetic variation modulates the susceptibility of pathological hypomethylation in a similar manner across most ICRs. The observed differences in the relative susceptibility of a subset of ICRs might reflect distinct regulatory mechanisms operational at these sites during development.

### Identification of QTLs determining susceptibility to locus-specific and widespread DNA hypomethylation

We leveraged the observed variation in DNA methylation levels at imprinted gene loci (**Fig.2B**) across DO nPSC lines to attempt to map quantitative trait loci (QTLs) for all ICRs (ICR-me QTLs) across 73 lines that passed additional QC criteria (see Methods). This resulted in the identification of six QTLs that reached statistical significance for at least one ICR (see Methods) (**Fig.2C**). Of note, none of these ICR-me QTLs were located on the same chromosome (Chr) as their putative target ICRs, indicating that each of the identified QTL regions functions in trans. Two ICR-me QTLs – located on Chr4 and Chr17 – are associated with multiple ICRs, with the Chr4 QTL showing the strongest association with ICRs from the *Grb10*, *Mest*, *Plagl1* and *Zsr1* loci and the Chr17 QTL associating strongest with ICRs at the *H19*, *Gtl2* promoter and *Zbdf2*_pat loci (**Fig.2C-E**). Several additional ICRs and non-imprinted genes showed strongly elevated LOD scores at these QTLs, suggesting that these regions harbor variants controlling the degree of DNA hypomethylation across multiple genomic target sites (**Fig.2C**).

The Chr4 QTL region (Chr4: 144,920,230-151,337,450) contains a large cluster of genes encoding KRAB zinc finger proteins (KRAB-ZFPs) (**Fig.2F**), which are rapidly evolving genes that play essential roles in silencing foreign DNA elements such as retrotransposons and endogenous retroviruses (Yang et al., 2017). Several prior genetic mapping studies have implicated this same genomic region in the regulation of DNA methylation, including as a modulator of transgene silencing (Engler et al., 1991) and of variably methylated IAP elements that can act as epi-alleles in mice (Bertozzi et al., 2020; Wolf et al., 2020). In addition, gene expression and chromatin accessibility QTL (caQTL) mapping in nPSCs from DO mice identified this region as a trans-eQTL/caQTL hotspot linked to changes in gene expression (115 affected genes) and chromatin accessibility (577 affected peaks) (Skelly et al., 2020). In contrast, to our knowledge, the region of Chr17 QTL (Chr17: 31,353,698-33,102,392) has not yet been implicated in regulating nPSC transcription or biology. Analysis of the Chr 17 QTL suggested several candidate genes that could contribute to the regulation of DNA methylation, including a cluster of genes encoding KRAB-ZFP proteins (*Zfp871*, *Zfp870*, *Zfp799*, *Zfp763*, *Zfp472*), the transcriptional co-activator *Brd4*, and *Wiz*, a known interaction partner of the repressive histone methyltransferases EHMT1 and EHMT2 (also known as GLP and G9a).

Of note, allelic effect analysis showed different contributions of specific DO founder strain haplotypes at the Chr4 and the Chr17 QTL (**Fig.2D, E**). In particular, B6J and 129 had opposite effects on DNA methylation changes mediated by the Chr17 QTL (**Fig.2E**), raising the possibility that this region might be involved in establishing the strain-specific DNA methylation stability observed in pure background nPSCs from these two strains. Among the genes encoded by the Chr17 QTL, transcriptomic analysis in B6J and 129 nPSCs revealed that *Wiz* is significantly higher expressed in B6J mESC cultured in both DMSO and iMEK conditions (**Fig.S2G**). *Wiz* was initially identified in an ENU screen for genes that modulate the rate of stochastic epigenetic silencing of an integrated reporter transgene (Daxinger et al., 2013) and the EHMT1/2 complex has been linked to the regulation of DNA methylation at several ICRs (Zhang et al., 2016). Further investigation will be needed to confirm the molecular reasons for the elevated *Wiz* expression levels in B6J mESCs and the potential role of WIZ in protecting ICR methylation stability. Our observations are consistent with strain-specific variants that affect the levels or activity of trans-acting epigenome regulators as major drivers of DNA methylation stability in mouse nPSCs.

## Discussion

The susceptibility of imprinted genes for dysregulation in mouse (Humpherys et al., 2001) and human (Kim et al., 2007) pluripotent cells and the associated risks (Greenberg and Bourc’his, 2015) have been well-established. More recently, efforts to establish naïve pluripotency in human cells have drawn additional attention to the issue of pathological DNA hypomethylation in these cells (Bar and Benvenisty, 2019). In contrast to the prevalent notion of the stochastic nature of imprint abnormalities (Humpherys et al., 2001), our results with inbred and genetically diverse nPSCs unambiguously demonstrate that genetic background contributes in a major way to variation in susceptibility to aberrant DNA hypomethylation during establishment and maintenance of naïve pluripotency. Based on QTL mapping conducted in DO nPSCs, we propose that genetic variants modulating the activity of specific trans-acting factors – including KRAB-ZFPs and candidate co-factors such as WIZ – represent additional critical variables contributing to epigenetic instability in nPSCs. Of note, while KRAB-ZFPs undergo rapid evolution (Yang et al., 2017), ZFP57 and ZFP445 have been reported to protect ICRs from demethylation in both mice and humans (Takahashi et al., 2019) (Juan and Bartolomei, 2019), raising the possibility that variants in these proteins may affect epigenome stability in nPSCs from both species.

The impact of genetic variation on imprint stability in cultured pluripotent cells has several significant ramifications. First, cell line- and locus-specific vulnerabilities complicate the identification of a universal media composition that can stabilize imprints across different cell lines. This is documented by the background-specific requirements of B6J and 129 nPSCs, with B6J nPSCs better tolerating de-methylating agents such as iMEK and AA and 129 essentially preserving imprints in serum-based culture conditions in the absence of additional compounds (Auerbach et al., 2000; Swanzey et al., 2020). Second, it may be possible to predict imprint stability in specific culture conditions based on the genetic variants present in a given nPSC line. Third, the targeted re-engineering of specific variants might enable stabilizing imprints in otherwise epigenetically unstable PSCs. In addition to these more practical considerations, the systematic identification and characterization of variants that impact imprint stability in nPSCs – for example, by analysis of larger panels of DO nPSC lines – might help to unravel the complex regulatory networks governing DNA methylation stability, with implications for a wide range of physiological and pathological processes.

## Acknowledgments

Subhashini Madhuranath for help with WB and Laurianne Scourzic for providing reprogramming virus. We are grateful to all Stadtfeld, Vierbuchen, and Apostolou lab members for their input on this project and feedback on this manuscript. M.S. was supported by grants from the NIH (R01GM145864), the Simons Foundation, the Tri-Institutional Stem Cell Initiative (Tri-SCI), and the Bohmfalk Charitable Trust. T.V. was supported by the Sloan Kettering Institute Josie Robertson Investigator Program, the Tri-Institutional Stem Cell Initiative (Tri-SCI), and the Sloan Kettering Institute Cancer Center Support Grant (NIH P30 CA008748). R.G. was supported via an NIH T32 training grant (T32 HD060600). J.G.M receives support from (R01GM145864, R01EB027918, R33HL151355, R01 HL166983).

## Author contributions

C.P. derived and characterized iPSCs from inbred mouse strains, isolated gDNA, and coordinated DNA methylation analysis by targeted bisulfite sequencing. R.A.G. led the effort to generate and characterize mESCs from DO mice, with assistance from S.S. and S.D, Y.F. performed in vitro fertilization and mESC derivation for Diversity Outbred mESC lines, Y.S. conducted QTL mapping, K.C conducted WB experiments and analyses and prepared samples for RNA-seq, E.E.S. derived inbred mESCs, Y.Z. analyzed DNA methylation data, T.M. conducted TLC, E.A. supervised data analysis and contributed to conceiving the study, A.P. analyzed RNA-seq data, R.K. supervised bioinformatic analysis, J.G.M. supervised QTL mapping, M.S. and T.V. conceived the study, supervised experiments, and wrote the manuscript with input from all authors.

---

**Figure S1.**
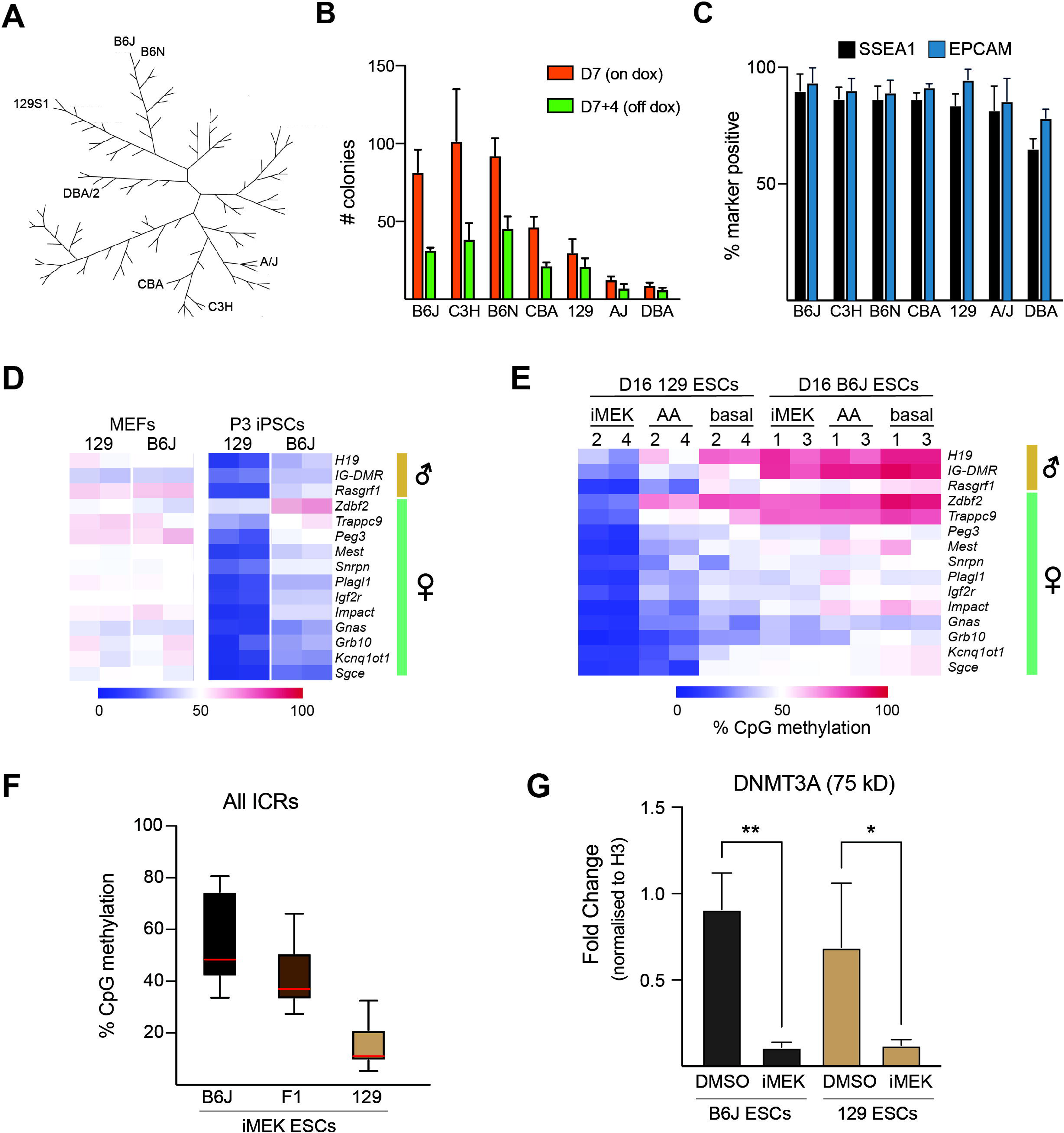
Genetic variation modulates DNA hypomethylation at ICRs in naive pluripotent stem cells from distinct inbred mouse strains. A. Phylogenetic tree of inbred mouse strains. Inbred strains used for reprogramming experiments are highlighted. Modified after (Petkov et al., 2004). B. Number of iPSC colonies obtained via reprogramming with nPSC morphology from each inbred mouse strain. Colony quantification was performed at two time points – D7: before withdrawal of doxycycline (viral OKSM expression active), D7+4: after doxycycline withdrawal (viral OKSM expression inactive). C. Quantification of pluripotency-associated surface marker expression (SSEA1, EpCAM) by flow cytometry in iPSCs from different inbred strains. D. DNA methylation levels at indicated ICRs in MEFs and iPSCs (passage 3) from 129 and B6J backgrounds. E. DNA methylation levels at ICRs in 129 and B6J mESCs after culture in indicated conditions. F. ICR methylation levels in B6J, 129, and F1 mESCs cultured in the presence of iMEK for 16 days. G. Quantification of DNMT3A protein levels in 129 and B6J mESCs after culture in iMEK or DMSO. Protein levels are normalized to histone H3 expression. * = p<0.05. and ** = p<0.01 with one-way ANOVA. N = 3 biological replicates.

**Figure S2.**
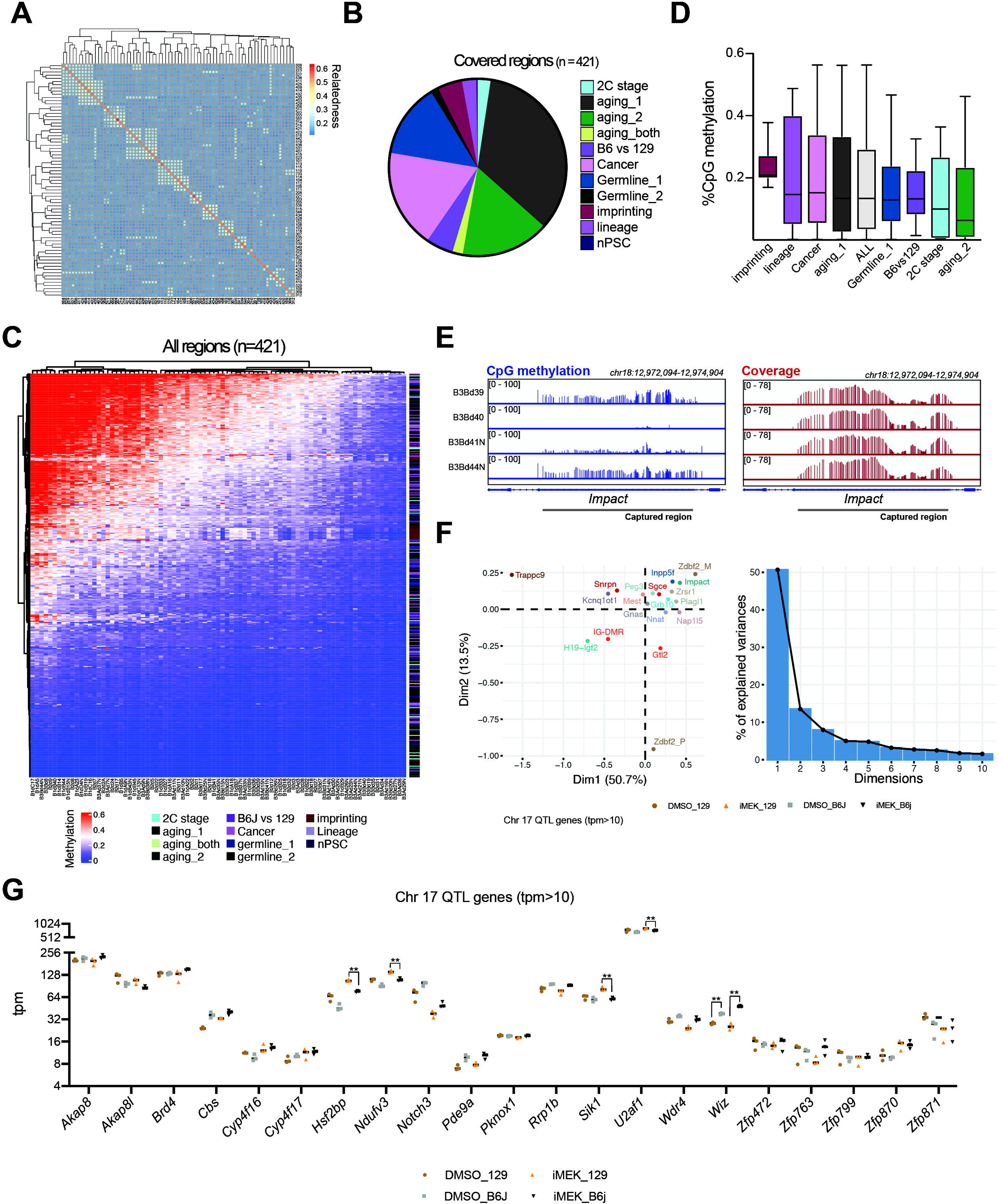
Identification of QTLs modulating pathological DNA hypomethylation at ICRs in naïve PSCs. A. Kinship diagram of the DO mESC lines (listed by GigaMUGA ID) used for QTL mapping. B. Gene category representation on capture array C. DNA methylation across all 421 covered regions in all DO mESCs D. DNA methylation levels in D0 mESCs at gene loci (mean and range) representing indicated gene categories. E. CpG methylation levels and coverage at *Impact* locus in four representative DO mESCs. F. PCA analysis of DNA methylation levels at ICRs G. RNA-seq expression levels of all genes encoded by the Chr17 QTL with tpm>10 in either B6J or 129 mESCs. ** = q<0.01 with multiple t-tests and Benjamini, Krieger, and Yekutieli corrections.

## Supplementary Tables

**Table S1: Table of pluripotent stem cell lines used in this study**

**Table S2: Table of genomic regions selected for targeted enrichment DNA methylation profiling**

## Resource availability

### Lead Contact

Requests for resources and reagents should be directed to and will be fulfilled by the lead contact, Matthias Stadtfeld.

## Materials availability

Cell lines generated in this study are available upon request from the lead contact. Diversity Outbred mESC lines are described in (Glenn et al., 2024) and can be requested from Thomas Vierbuchen.

## Experimental model and subject details

### Cell lines

Details of the cell lines used in this study are presented in **Table S1**.

## Method details

### Mouse embryonic fibroblast derivation and culture

Breeding age males and females from inbred strains 129S1/SVlmJ (129; JAX_002448), C57BL6/J (B6J; JAX_000664), C57BL6/NJ (B6N; JAX_005304), CBA/J (CBA; JAX_00656), DBA/2J (DBA; JAX_00671), C3H/HeJ (C3H; JAX_000659) and A/J (AJ; JAX_000646) were ordered and set up for mating. Embryos from visibly pregnant females were isolated between E14.5 and E16.5 and dissected from the uterus. After removal of the head and internal organs, the remainder of the embryo was placed into two drops of trypsin-EDTA (0.25%), finely minced using scalpels, and incubated at 37 degrees centigrade for five minutes. Then, embryos were dissociated in MEF media (DMEM with 10% fetal bovine serum, non-essential amino acids, GlutaMAX, pen/strep and 2-Mercaptoethanol) using a 10ml stripette, transferred into T75 flasks (one flask/embryo) and incubated in a 4% O_2_ incubator. Once fibroblast outgrowths covered most of the culture surface, cells were harvested by trypsinization, passed through a 40 μM cell strainer to remove remaining undissociated tissue, and frozen down or directly used for reprogramming experiments. A small number of cells was used for genomic DNA prep using a home-made lysis buffer consisting of 100mM Tris-HCl (pH8), 5mM EDTA, 200mM NaCl, 0.2% SDS, and 1 mg/ml Protein K in water. PCR with the primer pair JarifF (CTGAAGCTTTTGGCTTTGAG) and JaridR (CCACTGCCAAATTCTTTGG) was used to determine the sex of MEFs.

### Generation of induced pluripotent stem cells from mouse embryonic fibroblasts

Male MEFs at P1 were seeded in MEF media at 40% confluency onto 48-well plates (n=3 three wells MEFs from different embryos for each strain) and transduced with dox-inducible polycistronic STEMCCA lentivirus encoding OCT4, KLF4, SOX2 and MYC and a constitutive lentivirus expression the reverse tetracycline-dependent transactivator (rtTA) (Sommer et al., 2009). The day of transduction, media was changed to mouse ESC medium (KO-DMEM with 15% FBS, L-Glutamine, penicillin-streptomycin, non-essential amino acids, 2-Mercaptoethanol, and 1000 U/ml LIF) supplemented with doxycycline (dox; 1 μg/ml), L-ascorbic acid (AA; 50 μg/ml), TGFβ RI Kinase Inhibitor II (iALK5; 250nM) and CHIR99021 (CHIR; 3μM). Fresh reprogramming media was added daily until day 7 when colonies with ESC-like morphology were scored, and culture continued in mESC ESC media supplemented with AA but without dox, iALK5, and CHIR. Three to four days later, colonies that maintained ESC-like morphology were counted, and wells were passaged entirely onto 6-well plates. Culture in mESC supplemented with AA was maintained for three passages. At P3, a subset of cells was used for genomic DNA extraction, while the remaining cells were cultured until P15 in base mESC media before being used for DNA extraction.

### Mouse ESC derivation and culture conditions

ESC lines from inbred mouse strains were derived from individual E3.5 blastocysts as previously described (Czechanski et al., 2014). In brief, blastocysts were isolated and cultured in KnockOut DMEM (GIBCO) supplemented with 15% KnockOut Serum Replacement (GIBCO), 1% FBS, L-Glutamine, sodium pyruvate, penicillin, streptomycin, non-essential amino acids, 2-Mercaptoethanol, 1000 U/ml LIF, 1 μM PD03259010 and 3 μM CHIR99021 on a layer of Mitomycin-C-treated feeders for four days. Media was then changed to mESC medium on Mitomycin-C-treated feeder cells. If applicable, l-Ascorbic acid (50 μg/ml) or PD03259010 (1 μM) was added. ESC lines from DO mice were generated via a pooled in vitro fertilization procedure. The DO ESC lines described here are part of a larger panel of DO ESCs we generated (Glenn et al., 2024). Critical experimental details are reproduced here for the convenience of the reader. Briefly, In-vitro fertilization (IVF) was performed as described previously (Nakagata, 2011). On the day of IVF, sperm from fertility-tested DO male mice was collected from the cauda epididymis and pre-incubated in a drop of sperm medium (TYH-MBCD medium (prepared in-house) or Fertiup (CosmoBio, catalog #KYD-002-05-EX) for one hour. Cumulus masses containing oocytes were collected from the oviducts of superovulated females and placed in a drop of IVF medium (human tubal fluid (HTF) medium (prepared in-house) or Cook IVF medium (Cook Medical, catalog # K-RVFE). One to 4 uL of sperm was added to each IVF drop. For each batch of IVF, a non-sibling cohort of 10-15 males and 10-15 females was used. Sperm harvested from 5 males was pooled into 4-6 sperm incubation drops in different combinations. Likewise, oocytes from 5 females were collected into 4-to 6 IVF drops and then inseminated with distinct sperm pools. Three to 4 hours after insemination, eggs were rinsed to remove excess sperm and cumulus cells and cultured further. The following day, 2-cell stage embryos were collected and transferred to KSOM embryo culture medium (Embryo Max KSOM, Millipore-Sigma, catalog # MR-121-D) and further cultured for an additional 2-3 days to hatching/hatched blastocysts. Blastocysts were transferred to DO ESC culture medium (Cellartis 3i mES/iPSC medium, Takara Bio, catalog # 1181722446) in a cell culture dish covered with mouse embryonic fibroblast (MEF) feeder cells.

DO ESC derivation was performed as described by (Kiyonari et al., 2010). Briefly, outgrowths of inner cell mass from seeded blastocysts cultured in 3i medium, typically for 7-10 days, were manually picked. Cell clumps were dissociated with trypsin (0.25% Trypsin-EDTA, Gibco, catalog # 25200-056) and replated into MEF feeder-coated cell culture wells (typically, a 96-well plate). Cells were gradually expanded into larger cell culture wells, and the initial frozen stock vials were prepared from 6-well cell culture plates (typically, at passages 3 to 4). A few remaining cells were re-plated onto gelatin-coated, feeder-free cell culture wells/dishes for further expansion and genomic DNA preparation.

Primary stocks of Diversity Outbred mouse ESC lines (banked at passages 3-4) were thawed and expanded one additional passage on gelatin-coated dishes with irradiated mouse embryonic fibroblast feeder cells using serum/LIF + 2i media composed of DMEM (high glucose, GlutaMAX, HEPES), 1% nonessential amino acids, 1% sodium pyruvate, 1% penicillin-streptomycin, 0.1% 2-mercaptoethanol, 10% fetal bovine serum, 1000 U/ml ESGRO LIF, 3 µM CHIR99201, and 1 µM PD0325901. Media was changed daily, and ESCs were passaged upon 70% confluence at a 1:6 ratio using Accutase. DO ESC secondary stocks were banked at p4-5, and these stocks were thawed and used for DNA methylation profiling experiments. Mycoplasma testing was performed using the Mycoplasma PCR Detection Kit (Abm, Cat # G238), and all lines tested negative.

### Genomic DNA isolation for bisulfite sequencing and TLC

Genomic DNA was isolated using proteinase K in lysis buffer, pH 8 (100 mM Tris-HCl, 5 mM EDTA, 0.2% SDS, 200 mM NaCl), followed by isopropanol precipitation, ethanol wash, and reconstitution in TE buffer, pH 7.5 (10 mM Tris-HCl, 1 mM EDTA).

### Bisulfite sequencing

EpigenDx conducted bisulfite sequencing for the following ICR regions (assay and coordinates in mm10 in parentheses): H19 (ADS445; chr7:142,580,660-142,580,840), IG-DMR (ADS1452S; chr12:109,528,348-109,528,522), Rasgrf1 (ADS936R; chr9:89,879,735-89,879,870), Zdbf2 (ASY1814; chr1:63,263,595-63,263,743), Trappc9 (ASY1807; chr15:72,809,239-72,809,353), Peg3 (ASY1691; chr7:6,730,482-6,730,626), Mest (ADS915B; chr6:30,736,975-30,737,097), Snrpn (ASY1694; chr7:60,004,822-60,004,926), Plagl1 (ADS191; chr10:13,091,014-13,091,150), Igf2r (ASY1809R; chr17:12,742,271-12,742,174), Impact (ASY1812; chr18:12,974,427-12,974,544), Gnas (ASY1679; chr2:174,295,026-174,295,059), Grb10 (ASY1802; chr11:12,026,650), Kcnq1ot1 (ADS913; chr7:143,295,167-143,295,375) and Sgce (ASY1684; chr6:4,746,936-4,747,087).

### Analysis of 5mC levels using thin-layer chromatography

Nuclei were prepared by resuspending cells in 1 ml NPB (240 mM sucrose, 7.5 mM Tris, pH 7.5, 3.75 mM MgCl_2_, 0.75% Triton-X-100, with 100 μg RNAseA/ml (Qiagen 158922) and placing on ice for 20 minutes. Cells were spun at 1300 g for 15 min, 4 C, then washed once in NPB. Nuclei were lysed in 650 μl of 1X LB ((10 mM Tris, pH 8.0, 300 mM NaAcetate, pH 7.2, 0.5% SDS, 5 mM EDTA, 100 mg RNAseA/ml and 300 μg/ml Proteinase K (Roche 3115801001)) and incubated overnight at 55 C. An extra 300 μg/ml Proteinase K was added in the morning, and the samples were left at 55 C for 5 hours. Samples were extracted with equal volumes of phenol, phenol:chloroform: isoamyl alcohol (25:24:1), and chloroform: isoamyl alcohol (24:1), and then precipitated with two volumes of ethanol. Genomic DNA was washed twice with 1 ml of 70% EtOH, dried, resuspended in 10 mM Tris, 0.1 mM EDTA, pH 8.0, and resuspended overnight at 32 C.

2 μg of genomic DNA was digested with 100 units of Taq1-v2 (NEB R0149S) and 100 μg of RNaseA overnight. An extra 100 units of restriction enzyme was added in the morning, and incubations continued for 6 hours. Ten units of calf intestinal phosphatase (CIP) (NEB M0290L) were added and incubated for 1 hour at 37 C. DNA was purified using Qiaquick Nucleotide Removal Kit (Qiagen 28306) as per the manufacturer’s instructions. 400 ng of eluted DNA fragments were end-labeled with T4 Polynucleotide Kinase (T4 PNK) (NEB M0201L) and 10 μCi of [*γ*−^32^P]-ATP for 1 hour at 37 C. Labeled fragments were precipitated by the addition of 30 μg of linear polyacrylamide, 1/10 volume of 3 M NaAcetate, pH 7.2 and 2.5 volumes of ethanol at left at -80 C for 1 hour. Samples were spun at 14,000 rpm for 20 minutes at 4 C and washed twice with 70% EtOH at 25 C. Pellets were resuspended in 30 mM Tris, pH 8.9, 15 mM MgCl_2_, 2 mM CaCl, with 10 μg of DNaseI (Worthington LS006331) and 10 μg SVPD (Worthington LS003926) and incubated for 3 hours at 37 C. 3 μl was spotted on cellulose TLC plates (20 cm x 20 cm, Merck) and developed in isobutyric acid: H_2_0: NH_3_ (66:20:1). Plates were analyzed by phosphorimager scanning using Phosphorimager Storm 860 scanner software. The low-level labeling of other nucleotides reflects DNA shearing or contaminating endonucleolytic activity.

### Flow cytometry

For quantification of surface marker expression, trypsinized PSCs were stained with antibodies against EpCAM/CD326 (G8.8, Thermo) and SSEA1/CD15 (MC-480; Thermo), acquired on a FACSCanto (BD Biosciences) and analyzed with FlowJo software (Tree Star Inc.).

### Sample preparation for RNA-seq and Western blotting

B6J and 129 ES cells at passage 8 were treated in triplicates with either DMSO or at 1μM iMEK (PD 0325901; Tocris #4192) from day 0 to day 5 daily. Cells were collected by trypsinization at the end of 5 days, and samples were prepped separately for RNA and protein isolation.

### Western blotting

Cells were washed with PBS -/- and harvested with Trypsin. Nuclear lysates were prepared using the NE-PER kit (Thermo 78833) according to the manufacturer’s instructions. Protein concentration was measured using Bradford Reagent (BioRad #5000006), and samples were boiled in Laemmli Sample Buffer and run using Invitrogen NuPAGE 4-12% Bis-Tris precast gel (NP0322BOX). Western blots were performed using the following antibodies: anti-DNMT3A (Active Motif # 39206, anti-pERK1/2 (CST #9101), and anti-histone H3 (Abcam #ab1791).

### RNA-seq

Total RNA was extracted using TRIzol (Invitrogen 15596018) and purified with the RNA Clean and Concentrator kit (Zymo Research ZR1014). Following the manufacturer’s instructions, one ug of RNA was used to make libraries using the TruSeq Stranded mRNA Library Prep Kit (Illumina# 20020595). Libraries were sequenced on the NovaSeq 6000 using an S4 flow cell at PE 2X100 at the Genomics Core of Weill Cornell Medicine.

### SNP Genotyping of Diversity Outbred ESC lines

DNA was extracted from ESCs grown on feeders for 1-2 passages (5e5 to 1e6 total cells/sample). DNA was extracted using the Qiagen DNeasy Blood & Tissue Kit (#69504), followed by the Zymo DNA Clean and Concentrator. Double-stranded DNA content was measured using a NanoDrop OneC Spectrophotometer (Thermo Fisher, #13-400-518). Extracted DNA was aliquoted in a 96-well qPCR plate in at least 10 uL of volume at greater than 20 ng/uL concentration in groups of at least 24 samples. Samples were shipped to Neogen Genomics (Lincoln, NE) on dry ice overnight for sequencing via the GigaMUGA genotyping array on the Illumina Infinium platform (Morgan et al., 2015).

### DO cell culture/sample collection for hypomethylation experiments

DO ESCs were thawed at passages 4-5 on gelatin-coated dishes with irradiated feeder cells in Serum/LIF +2i media and cultured for another two to four days. To avoid feeder contamination, ESCs were harvested by detaching colonies using collagenase IV (500 U/mL). Cells were collected and centrifuged at 300xg for 3 minutes at room temperature and stored at -20C before being shipped to SAMPLED (Piscataway, NJ) for processing using Twist NGS Methylation Detection Workflow.

### Design of targeted capture panel

A Custom Twist Methylome Panel (Twist Bioscience) was designed to perform targeted enrichment for genomic regions chosen based on their known functions as ICRs (Dahlet et al., 2020; Swanzey et al., 2020) or association with germline development (Mochizuki et al., 2021), mouse pre-implantation development (Hu et al., 2020), post-implantation development (Dahlet et al., 2020), aging (Meer et al., 2018; Stubbs et al., 2017), or cancer (Brady et al., 2021). Coordinates of all targeted regions are provided in **Table S2.**

### DNA methylation profiling and targeted enrichment

DNA methylation profiling was performed using the Twist NGS Methylation Detection Workflow. Briefly, the NEBNext Enzymatic Methyl-Seq Library Preparation protocol (https://www.twistbioscience.com/resources/protocol/neb-next-enzymatic-methyl-seq-library-preparation-protocol) was used to prepare genome-wide DNA methylation libraries starting from 200 ng gDNA per sample. EM-seq libraries (187.5 ng/sample) were then used for targeted enrichment via the Twist Targeted Methylation Sequencing Protocol (https://www.twistbioscience.com/resources/protocol/twist-targeted-methylation-sequencing-protocol). Hybridization was performed at 60 C, eight samples were multiplexed for each hybridization reaction, and 15 cycles of PCR were performed post-capture. Libraries were sequenced using the MiSeq platform (2 × 150 bp reads). The Twist NGS Methylation Detection Workflow was performed by SAMPLED (Piscataway, NJ).

## DATA ANALYSIS

### DNA methylation data processing and quantification

Bismark pipeline (Krueger and Andrews, 2011) was adopted to map DNA methylation sequencing reads and determine cytosine methylation states. Using Trim Galore v0.6.4 (https://github.com/FelixKrueger/TrimGalore), raw reads with low-quality (less than 20) and adapter sequences were removed (Martin, 2011). The trimmed sequence reads were C(G) to T(A) converted and mapped to similarly converted reference mouse genome (mm10) using default Bowtie 2 (Langmead and Salzberg, 2012) settings implemented by Bismark. Duplicated reads were discarded. The remaining alignments were then used for cytosine methylation calling by Bismark methylation extractor. Furthermore, the CpG sites with read coverage less than 10 were removed from the downstream analyses. In this study, we focus on the targeted regions. The full count matrix was mapped to the targeted regions using the function getCoverage() with the regions from R package bsseq (v1.26.0) (Hansen et al., 2012). Average coverage per region was calculated by dividing the total region coverage by the number of captured Cs. The methylation level per region was represented by the average methylation level of all CpGs in the respective region. Average methylation levels were measured for 421 regions, including 19 ICRs or DMRs at imprinted loci. These 19 regions were considered for QTL mapping, along with a 20th composite trait calculated as the within-line average of the methylation levels for all 421 regions.

### SNP genotyping data analysis

The sample sequences were processed using the R/qtl2 package to encode the SNP genotypes (Broman et al., 2019b). Subsequently, we performed a kinship analysis to compare the genotype probabilities between all DO lines used in this study and define the degree of relatedness between lines from the same IVF preparation as adapted from (Manichaikul et al., 2009) (https://smcclatchy.github.io/mapping/). The following were computed for each diversity outbred mouse ***(see above methods***): X chromosome heterozygosity, number of crossovers, proportion missing data, and proportion heterozygous sites (Broman et al., 2019a). The Y chromosome intensity per mouse was calculated by averaging the average Y chromosome microarray signal across all SNPs.

For the QTL mapping analyses, mice were included if they satisfied the following Quality Control criteria [0]: Either have X chromosome heterozygosity < 0.1 and Y chromosome intensity > 0.2 (i.e., mice considered to be male), between 600 and 900 crossovers, less than or equal to 2% proportion missing genotypes, and proportion heterozygosity between 0.75 and 0.9. In addition, in cases of duplicates, only one was retained. Finally, plots of the B allele frequency and the log R ratio per sample were generated using the karyoploteR package (Gel and Serra, 2017) to assess for chromosomal abnormalities; samples with a B allele frequency that deviates from 0.5 at any chromosome were excluded. These total criteria resulted in 73 cell lines. We prepared genotype data using the pipeline (Broman et al., 2019b) with reference genome assembly GRCm38(mm10), obtaining 105,200 SNPs with mappable rsIDs. Genotype quality control measures were applied, including filters for Minor Allele Frequency (MAF) of 0.125, missingness of 0.02, and Hardy-Weinberg Equilibrium (HWE) with a threshold of 1e-6, where following these filters, the final genotype dataset analyzed consisted of 84,403 SNPs.

### RNA-seq Analysis

Paired-end read alignment to the mouse genome (mm10 version) was performed with STAR aligner (version 2.7.10)□ with the default setting. Samtools (version 1.13)(Li et al., 2009) were used for filtering and sorting aligned reads before annotation to “mm10.GRCm38.95.gtf” gene version with htseq-count (0.6.1p1 version)(Anders et al., 2015). Only protein-coding and long-non-coding RNA transcripts were used for annotation, and downstream differential expression analysis was performed using the R package DESeq2 (Love et al., 2014). Genes with p-adjusted value <0.01 and fold change difference of 2 were considered differentially expressed.

### QTL mapping analyses

QTL mapping was performed for the 20 regions, considering the 73 ESC lines that passed the quality control criteria. The 20 traits were found to be approximately normal, where we found repeating mapping analyses after normal transformations did not appreciably change the outcome. Given the complexity of the DO design, we considered two QTL mapping analyses that employed slightly different but complementary modeling approaches: R/qtl2 (Broman et al., 2019b) and GEMMA (Zhou and Stephens, 2012). We implemented both to apply a mixed model with an additive (one-degree of freedom) bi-allelic coding of the focal marker, such that the differences between the two approaches are: R/qtl2 includes imputation of the bi-allelic state considering the eight founder genomes and a relatedness matrix for the random effect is calculated using the “leave-one-chromosome-out” (LOCO) method (Yang et al., 2014), whereas GEMMA analysis is performed on only measured genotypes and the relatedness matrix is calculated as the covariance of the centered genotypes (Zhou and Stephens, 2012). The implementation of the R/qtl2 analysis followed the user guide (Broman et al., 2019b), including write_control_file() with 40, 42, or 46 generations as appropriate per line, to create founder_geno_file, gmap_file, and pmap_file, insert_pseudomarkers() to insert pseudo markers into the genetic map, calc_genoprob() to calculate the QTL genotype probabilities, conversion of the pr to apr using alleleprob(), LOCO method to calculate the relatedness matrix, performing the genome scan with scan1(), and finding LOD peaks with find_peaks(out, map, threshold=5, drop=1.5). For the GEMMA analysis, the option -gk 1 was used for the relatedness matrix, both Wald and likelihood ratio tests were calculated, and Quantile-Quantile (QQ) plots of the p-values were used to assess the appropriateness of the model fit.

To assess significance, we applied family-wise error rate (FWER) corrections, where for R/qtl2 analyses, we used both a permutation approach (as implemented in R/qtl2) and Benjamini-Hochberg adjusted p-value (Benjamini et al., 2009) for the p-values obtained by converting the LOD scores at the measured genotypes to Likelihood Ratio Test statistics. For the GEMMA analyses, we applied the Benjamini-Hochberg adjusted p-value procedure. Given the smaller sample size of this study (73) compared to other DO QTL mapping studies that employed considerably larger samples (e.g., ∼300-500), (Keller et al., 2019; Linke et al., 2020; Price et al., 2023) we adopted a relatively liberal cutoff of a FWER <0.1 attained in either analysis to indicate a QTL. Similar to other studies, we also consider additional “suggestive” QTL that did not reach this FWER cutoff, (Keller et al., 2019; Linke et al., 2020; Price et al., 2023) whereas implemented in (Linke et al., 2020), we considered a FWER <0.2 attained in either study to indicate suggestive QTL.

### Quantification and statistical analyses

Statistical analysis of flow cytometry and IF data was done in PRISM 9 (GraphPad), with specific tests and corrections applied as indicated in the respective figure legends.

## DATA AVAILABILITY

Source data are available in the Gene Expression Omnibus under the accession numbers GSE268906 (EM-seq) and GSE267262 (RNA-seq).

